# Ultrasensitive single-cell proteomics workflow identifies >1000 protein groups per mammalian cell

**DOI:** 10.1101/2020.06.03.132449

**Authors:** Yongzheng Cong, Khatereh Motamedchaboki, Santosh A. Misal, Yiran Liang, Amanda J. Guise, Thy Truong, Romain Huguet, Edward D. Plowey, Ying Zhu, Daniel Lopez-Ferrer, Ryan T. Kelly

## Abstract

We report on the combination of nanodroplet sample preparation, ultra-low-flow nanoLC, high-field asymmetric ion mobility spectrometry (FAIMS), and the latest-generation Orbitrap Eclipse Tribrid mass spectrometer for greatly improved single-cell proteome profiling. FAIMS effectively filtered out singly charged ions for more effective MS analysis of multiply charged peptides, resulting in an average of 1056 protein groups identified from single HeLa cells without MS1-level feature matching. This is 2.3 times more identifications than without FAIMS and a far greater level of proteome coverage for single mammalian cells than has been previously reported for a label-free study. Differential analysis of single microdissected motor neurons and interneurons from human spinal tissue indicated a similar level of proteome coverage, and the two subpopulations of cells were readily differentiated based on single-cell label-free quantification.

---

Mass spectrometry (MS)-based proteome profiling provides insights into biological function and dysfunction that are unavailable through genomic or transcriptomic measurements.^[1]^ Extending proteomic analysis to single cells and other low-input samples sheds additional light on the roles of various cell types contributing to normal and disease processes and can yield spatial information for tissue mapping and characterization of the microenvironment.^[2]^ Given the absence of amplification techniques for proteins, every aspect of the analytical method must be carefully optimized to bring more protein species above detection limits to provide a more comprehensive view of protein expression, ideally extending to thousands of proteins per cell. These optimization efforts span the entire proteomics workflow, from cell isolation and sample preparation to MS measurement and data processing, and it is the combination of these advances that has made single-cell proteomics possible. For example, efforts to miniaturize sample preparation to nanoliter volumes using, e.g., nanoPOTS,^[3]^ the oil-air-droplet (OAD) chip^[4]^ or the integrated proteome analysis device (iPAD),^[5]^ have effectively reduced adsorptive losses and increased sample concentrations for more efficient protein digestion for single cells and other trace samples. Ultrasensitive separations have been realized by reducing total flow rates to the low-nanoliter-per-minute range using capillary electrophoresis^[6]^ or narrow-bore liquid chromatography (LC) with either open tubular^[7]^ or packed columns,^[8]^ providing reduced solvent contamination and improved ionization efficiency at the electrospray source.

Using a combination of nanoPOTS sample preparation, nanoLC separations operated at 20 nL/min and the Orbitrap Eclipse Tribrid mass spectrometer, we recently identified an average of 362 and 874 protein groups from single HeLa cells^[8b]^ without and with the Match Between Runs (MBR) algorithm of MaxQuant, respectively, which was the highest level of coverage reported for a label-free analysis of single mammalian cells. While this coverage is sufficient to differentiate between distinct cell types^[9]^ and illuminate processes involving high-abundance proteins,^[10]^ current methods are blind to expression patterns of lower abundance proteins that fall below detection limits. TMT-based approaches that incorporate a boosting channel^[11]^ can increase single-cell proteome coverage, but the quantitative accuracy is currently compromised by batch effects, ratio compression and the ‘carrier proteome effect’.^[12]^ Additional sensitivity gains achieved by further optimizing the analytical workflow are expected to improve both label-free and isobaric labeling methods.

During LC-MS analysis, tryptic peptides may be present as singly charged or multiply charged ions^[13]^ while most contaminating species and solvent clusters are singly charged. To increase MS/MS sequencing efficiency, only multiply charged species are typically selected for fragmentation, yet the presence of these +1 species in the MS1 scan increase spectral complexity and singly charged ions may still be co-isolated for fragmentation along with selected peptides, interfering with identification. More importantly, in the case of ion trapping instruments such as the Orbitrap, singly charged ions may occupy a significant portion of the trap capacity, effectively reducing sensitivity for multiply charged species and limiting those selected for fragmentation. The proportion of the ion population comprising singly charged species increases as sample size decreases, while solvent contributions remain relatively constant, so the presence of +1 ions is likely more detrimental for trace samples.

High field asymmetric ion mobility spectrometry (FAIMS)^[14]^ is a gas-phase separation technique in which the an asymmetric electric field is used to disperse ions and selectively filter ion populations by varying the compensation voltage (CV). Importantly, FAIMS can selectively remove +1 ions while broadly transmitting multiply charged peptides,^[15]^ which should be especially beneficial for single-cell proteomic analysis. However, some signal attenuation of selected ions occurs due to imperfect transmission through FAIMS devices, so it is necessary to determine whether the benefits of FAIMS overcome any detrimental decrease in signal. Here we evaluate the use of FAIMS for single-cell proteome profiling as depicted in Scheme 1. Single cells were isolated by capillary-based micromanipulation or laser capture microdissection (LCM) and processed in ~200-nL droplets in a nanoPOTS chip.^[3]^ Peptides were separated using a 20-μm-i.d. home-packed nanoLC column,[8b] ionized at a chemically etched nanospray emitter^[16]^ and then fractionated using the FAIMS Pro interface (Thermo, Waltham, MA) for selective removal of singly charged species and transmission of multiply charged peptides to the Thermo Scientific Orbitrap Eclipse Tribrid MS. Raw data were processed using Proteome Discoverer Software 2.4 (Thermo) or MaxQuant version 1.6.7.0.^[17]^

**Scheme 1.**
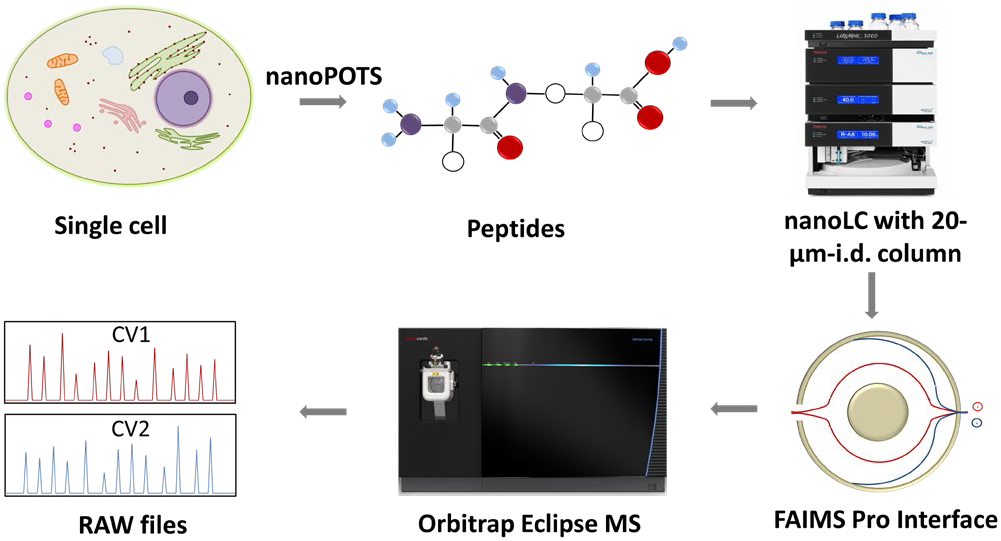
Single-cell proteomics workflow. Proteins from a single cell are extracted and digested, the resulting tryptic peptides are separated using a narrow-bore nanoLC column and ionized at an etched electrospray emitter. Singly charged ions are filtered using the FAIMS Pro interface and transmitted ions are detected using the Orbitrap Eclipse Tribrid MS.

MS acquisition and FAIMS settings were evaluated using 0.5 ng aliquots of commercial HeLa protein digest standard equivalent to 2–3 cells. Proteome coverage increased by ~30% when using the ion trap (IT) instead of the Orbitrap (OT) for MS2 (Figure 1A and Table S1), which is attributed to the higher sensitivity of the ion trap. Proteome coverage was ~12% higher when using HCD fragmentation rather than CID (Figure 1A). We also evaluated whether scanning between 2 or 3 compensation voltage (CV) values would provide greater coverage. Under the current conditions of a 120 min LC gradient, a 2 CV method (−55 V and - 70 V) provided ~10% greater coverage than a 3 CV method (−55 V, −70 V and −85 V) (Figure 1A). The 2 CV method with HCD fragmentation and ion trap detection was thus selected for single-cell studies. We note these settings will likely change with different sample loadings and LC gradients, etc.

**Figure 1.**
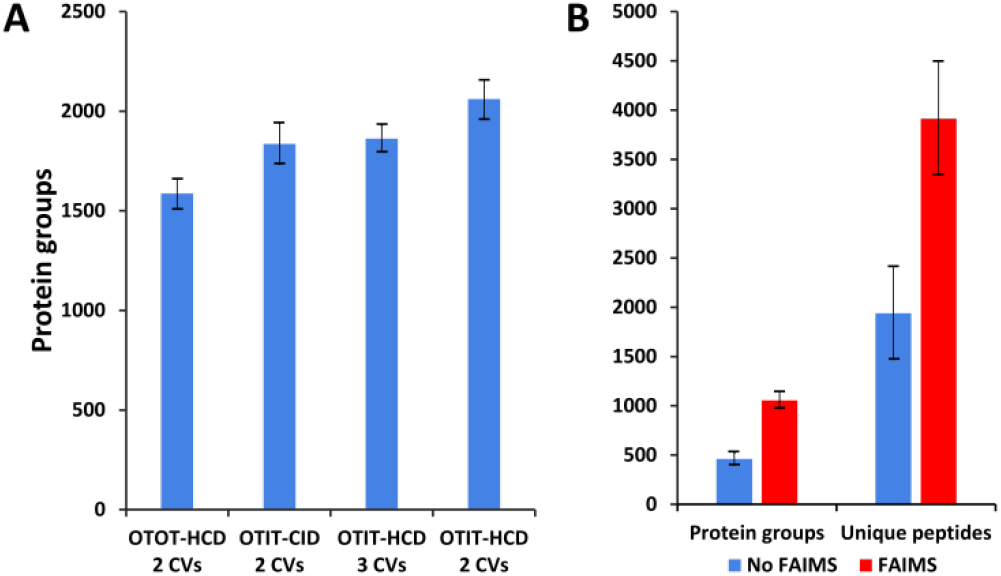
A) FAIMS method optimization using 0.5 ng aliquots of HeLa protein digest. Protein groups identified with different detection and fragmentation methods and using two or three FAIMS CVs. B) Protein groups and unique peptides identified from single HeLa cells with and without FAIMS. Error bars indicate standard deviations from 3 replicates.

Single HeLa cells were aspirated with 6 nL of supernatant, deposited into a nanoPOTS chip for sample preparation and analyzed as described above. Blank samples containing an equivalent volume of cell-free supernatant were analyzed in the same fashion to serve as a negative control. Mass spectra such as those shown in Figure 2 were compared for single HeLa cells with and without the FAIMS Pro interface. Without FAIMS, spectra were primarily composed of +1 ions, whereas FAIMS effectively filtered out most singly charged species. In evaluating proteome coverage for single HeLa cells, we identified on average 1056 protein groups from 3912 peptides when using FAIMS and Proteome Discoverer Software 2.4 with an FDR cutoff of <0.01 at both protein and peptide level, which represents a respective increase of 2.3 and 2.0-fold at the protein and peptide level compared to without FAIMS (Figure 1B and Table S2). We also evaluated proteome coverage using MaxQuant, which yielded an average of 683 and 1475 protein groups without and with MBR when including a matching library comprising 100 HeLa cells. While MaxQuant yielded fewer identifications without MBR relative to Proteome Discoverer, the level of proteome coverage was still 1.9 times greater than the 362 protein groups identified previously^8b]^ under identical analysis and database search conditions but without FAIMS.

**Figure 2.**
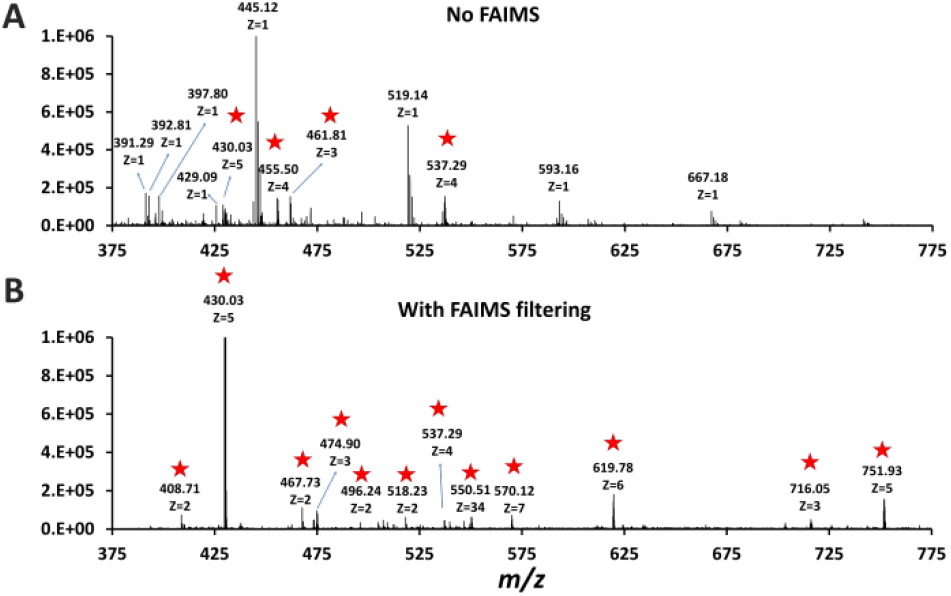
Representative mass spectra obtained without (A) and with (B) FAIMS filtering (CV −55 V). Peaks corresponding to multiply charged ions in both spectra are starrred.

To further explore platform performance on human post-mortem tissues and evaluate the ability of this platform to differentiate between closely related neuronal cell types, we applied our workflow to the analysis of single motor neurons (MNs) and interneurons (INs) excised by LCM from 12-μm-thick human spinal cord sections. Single-cell proteomic technologies represent an important platform for probing cell-type-specific perturbations, particularly in the context of human neurological diseases such as amyotrophic lateral sclerosis (ALS) and spinal muscular atrophy (SMA) characterized by selective vulnerability of motor neurons.^[18]^ An average of 1012 and 1085 protein groups were identified from single MNs (n=3) and INs (n=3), respectively, when applying a FDR cutoff of <0.01 and without MS1-level feature matching (Figure 3A). The identified protein groups from the two cell types had an overlap of 77% (Figure 3B) and were readily differentiated by principal component analysis (Figure 3C). Among the 1118 quantifiable protein groups (present with ≥2 unique peptides and in ≥50% of samples), 39 were significantly differentially abundant in MNs relative to INs (p<0.05, |Fold Difference|≥2) (Figure 3D, Supplemental Dataset II).

**Figure 3.**
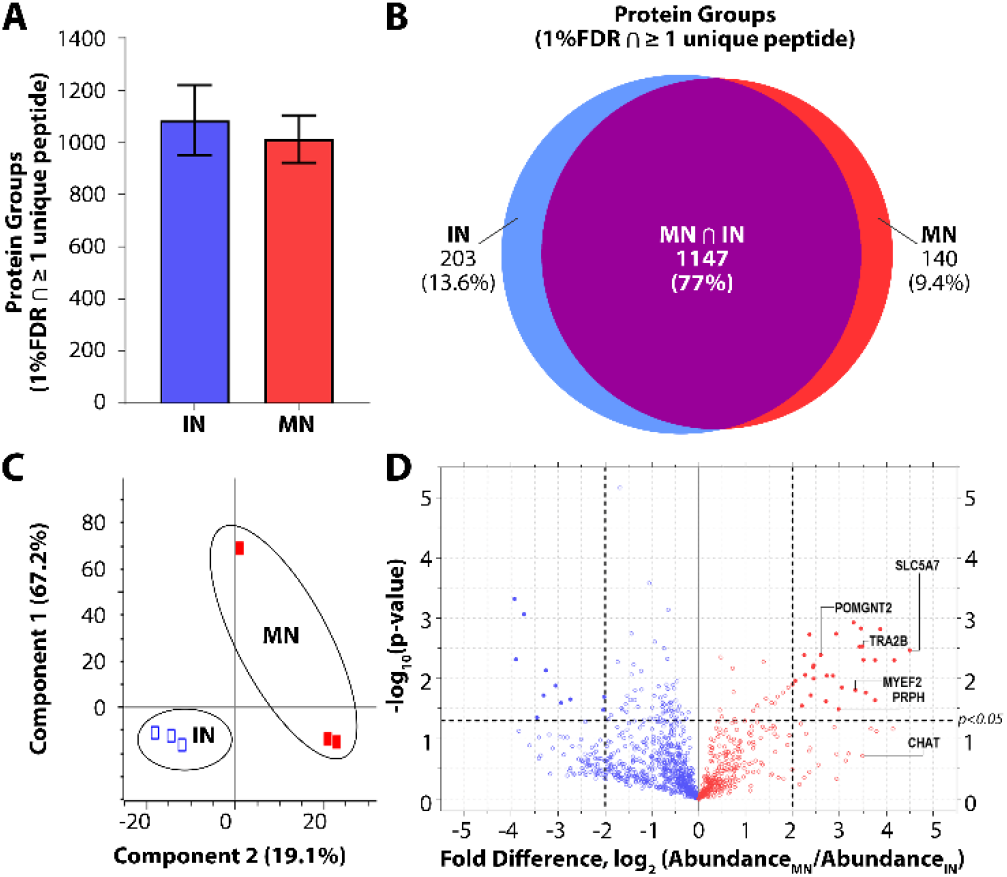
Single-cell proteomic interrogation of human spinal motor neurons and interneurons. A) Protein groups identified from single motor neurons (MNs) and interneurons (INs). B) Venn diagram indicating overlap of identified protein groups. C) Principle component analysis showing differentiation of the two neuronal subtypes based on 1118 quantified features. D) Volcano plot indicating significant differences in protein expression for quantifiable protein groups (p<0.05, |Fold Difference|≥2).

Gene ontology (GO) enrichment analyses on the subset of enriched-in-MN proteins revealed over-representation of proteins associated with RNA processing and alternative splicing, RNA metabolism, and post-transcriptional regulation of gene expression (Figure S1B). Consistent with established functions of MNs and mechanisms underlying MN-mediated stimulation of muscle fibers, we identified significant MN-enrichment of the high-affinity choline transporter protein SLC5A7 that mediates choline uptake in cholinergic neurons for rapid conversion into acetylcholine by choline o-acetyltransferase (CHAT) (Figure S1C). CHAT itself was approximately 10 times more abundant in MNs relative to INs (Supplemental Dataset II). Acetylcholine released by presynaptic MNs at the neuromuscular junction binds nicotinic receptors expressed on the post-synaptic membranes of muscle fibers.^[19]^ Furthermore, a number of proteins implicated in motor function and neuromuscular disease were represented among the subset of differentially abundant protein groups. These include RNA splicing machinery components such as TRA2B, the splicing factor that targets the survival of motor neuron (SMN) protein^[20]^ whose corresponding gene mutation causes SMA; MYEF2, a downstream splicing target of SMN;^[21]^ as well as multiple interactors of FUS and TARDBP (TDP43), whose corresponding gene mutations are causative for ALS (Figure S1C). FUS and TARDBP were detected in larger pools of MNs (5 cell pools) and in single ventral horns (*not shown*), but are likely below the limit of detection in single MNs. In addition to FUS and TARDBP, the intermediate filament protein peripherin (PRPH), important for axonal transport, is significantly more abundant in MNs relative to INs. Interestingly, mutations in PRPH are also associated with ALS,^[22]^ while overexpression of wild-type PRPH has been shown to result in selective motor axon degeneration in mice,^[23]^ suggesting regulation of PRPH expression is critical for motorneuron integrity. Increased PRPH abundance in spinal motorrelative to inter-neurons is consistent with previous observations that while PRPH is predominately expressed in the peripheral nervous system, it is also expressed in subsets of central nervous system neurons containing peripheral projections (e.g., MNs), and is reported to be highly expressed in lumbar spinal MNs at the mRNA level.^[24]^ Together, these data demonstrate the ability of unbiased proteomic profiling to (1) differentiate between neuronal subpopulations at the single-cell level and (2) identify differentially-expressed proteins and pathways relevant to cell-type-specific functions of MNs in health and disease.

The combination of nanoPOTS sample preparation, ultra-narrow-bore LC separation, Orbitrap Eclipse Tribrid mass spectrometer and the FAIMS Pro interface provides an unprecedented label-free proteome coverage of >1000 protein groups per mammalian cell using MS/MS identification alone. Additional gains will likely be realized by further miniaturizing sample preparation, pushing highly efficient nanoLC to lower flow rates, and optimizing FAIMS and MS acquisition settings. This platform promises to provide insights into cellular heterogeneity and enable the characterization of tissue microenvironments through in-depth mapping of protein expression in tissues with single-cell spatial resolution.

## Acknowledgements

Research reported in this publication was supported by the National Cancer Institute of the National Institutes of Health under award number R33 CA225248 and through a sponsored research agreement with Biogen, Inc. The content is solely the responsibility of the authors and does not necessarily represent the official views of the National Institutes of Health or Biogen, Inc.

## References

[1] T. E. Angel, U. K. Aryal, S. M. Hengel, E. S. Baker, R. T. Kelly, E. W. Robinson, R. D. Smith, Chem Soc Rev 2012, 41, 3912–3928.

[2]a aS. P. Couvillion, Y. Zhu, G. Nagy, J. N. Adkins, C. Ansong, R. S. Renslow, P. D. Piehowski, Y. M. Ibrahim, R. T. Kelly, T. O. Metz, Analyst 2019, 144, 794–807;

b bP. D. Piehowski, Y. Zhu, L. M. Bramer, K. G. Stratton, R. Zhao, D. J. Orton, R. J. Moore, J. Yuan, H. D. Mitchell, Y. Gao, B.-J. M. Webb-Robertson, S. K. Dey, R. T. Kelly, K. E. Burnum-Johnson, Nat Commun 2020, 11, 8;

[2]c cM. P. Snyder, S. Lin, A. Posgai, M. Atkinson, A. et al., Nature 2019, 574, 187–192.

[3] Y. Zhu, P. D. Piehowski, R. Zhao, J. Chen, Y. F. Shen, R. J. Moore, A. K. Shukla, V. A. Petyuk, M. Campbell-Thompson, C. E. Mathews, R. D. Smith, W. J. Qian, R. T. Kelly, Nat Commun 2018, 9, 882.

[4] Z. Y. Li, M. Huang, X. K. Wang, Y. Zhu, J. S. Li, C. C. L. Wong, Q. Fang, Anal Chem 2018, 90, 5430–5438.

[5] X. Shao, X. Wang, S. Guan, H. Lin, G. Yan, M. Gao, C. Deng, X. Zhang, Anal Chem 2018, 90, 14003–14010.

[6]a aL. L. Sun, G. J. Zhu, Y. M. Zhao, X. J. Yan, S. Mou, N. J. Dovichi, Angew Chem-Int Edit 2013, 52, 13661–13664;

b bC. Lombard-Banek, S. A. Moody, P. Nemes, Angew Chem-Int Edit 2016, 55, 2454–2458.

[7]a aS. Y. Li, B. D. Plouffe, A. M. Belov, S. Ray, X. Z. Wang, S. K. Murthy, B. L. Karger, A. R. Ivanov, Mol Cell Proteomics 2015, 14, 1672–1683;

b bP. L. Xiang, Y. Zhu, Y. Yang, Z. T. Zhao, S. M. Williams, R. J. Moore, R. T. Kelly, R. D. Smith, S. R. Liu, Anal Chem 2020, 92, 4711–4715.

[8]a aY. Zhu, R. Zhao, P. D. Piehowski, R. J. Moore, S. Lim, V. J. Orphan, L. Pasa-Tolic, W. J. Qian, R. D. Smith, R. T. Kelly, Int J Mass Spectrom 2018, 427, 4–10;

b bY. Cong, Y. Liang, K. Motamedchaboki, R. Huguet, T. Truong, R. Zhao, Y. Shen, D. Lopez-Ferrer, Y. Zhu, R. T. Kelly, Anal Chem 2020, 92, 2665–2671.

[9]a aY. Zhu, G. Clair, W. B. Chrisler, Y. F. Shen, R. Zhao, A. K. Shukla, R. J. Moore, R. S. Misra, G. S. Pryhuber, R. D. Smith, C. Ansong, R. T. Kelly, Angew Chem-Int Edit 2018, 57, 12370–12374;

b bM. W. Dou, Y. Zhu, A. Liyu, Y. R. Liang, J. Chen, P. D. Piehowski, K. R. Xu, R. Zhao, R. J. Moore, M. A. Atkinson, C. E. Mathews, W. J. Qian, R. T. Kelly, Chem Sci 2018, 9, 6944–6951;

c cY. Zhu, J. Podolak, R. Zhao, A. K. Shukla, R. J. Moore, G. V. Thomas, R. T. Kelly, Anal Chem 2018, 90, 11756–11759;

d dY. Zhu, M. W. Dou, P. D. Piehowski, Y. R. Liang, F. J. Wang, R. K. Chu, W. B. Chrisler, J. N. Smith, K. C. Schwarz, Y. F. Shen, A. K. Shukla, R. J. Moore, R. D. Smith, W. J. Qian, R. T. Kelly, Mol Cell Proteomics 2018, 17, 1864–1874.

[10] Y. Zhu, M. Scheibinger, D. C. Ellwanger, J. F. Krey, D. Choi, R. T. Kelly, S. Heller, P. G. Barr-Gillespie, Elife 2019, 8.

[11]a aB. Budnik, E. Levy, G. Harmange, N. Slavov, Genome Biol 2018, 19, 161;

b bM. W. Dou, G. Clair, C. F. Tsai, K. R. Xu, W. B. Chrisler, R. L. Sontag, R. Zhao, R. J. Moore, T. Liu, L. Pasa-Tolic, R. D. Smith, T. J. Shi, J. N. Adkins, W. J. Qian, R. T. Kelly, C. Ansong, Y. Zhu, Anal Chem 2019, 91, 13119–13127.

[12] T. K. Cheung, C.-Y. Lee, F. Bayer, A. McCoy, B. Kuster, C. M. Rose, in Proceedings of the 68th ASMS Conference on Mass Spectrometry and Allied Topics, 2020, MP 544.

[13] J. A. Loo, H. R. Udseth, R. D. Smith, Anal Biochem 1989, 179, 404–412.

[14] R. Guevremont, J Chromatogr A 2004, 1058, 3–19.

[15]a aD. A. Barnett, B. Ells, R. Guevremont, R. W. Purves, J Am Soc Mass Spectrom 2002, 13, 1282–1291;

b bD. B. Bekker-Jensen, A. Martinez-Val, S. Steigerwald, P. Ruther, K. L. Fort, T. N. Arrey, A. Harder, A. Makarov, J. V. Olsen, Mol Cell Proteomics 2020, 19, 716–729.

[16] R. T. Kelly, J. S. Page, Q. Z. Luo, R. J. Moore, D. J. Orton, K. Q. Tang, R. D. Smith, Anal Chem 2006, 78, 7796–7801.

[17] S. Tyanova, T. Temu, J. Cox, Nat Protocols 2016, 11, 2301–2319.

[18]a aH. J. Fu, J. Hardy, K. E. Duff, Nat Neurosci 2018, 21, 1350–1358;

b bM. Bowerman, L. M. Murray, F. Scamps, B. L. Schneider, R. Kothary, C. Raoul, European J Med Genetics 2018, 61, 685–698.

[19] C. R. Slater, Int Journal Mol Sci 2017, 18.

[20]a aY. Hofmann, C. L. Lorson, S. Stamm, E. J. Androphy, B. Wirth, Proc Nat Acad Sci USA 2000, 97, 9618–9623;

b bY. Hofmann, B. Wirth, Human Mol Genetics 2002, 11, 2037–2049.

[21] S. K. Custer, T. D. Gilson, H. X. Li, A. G. Todd, J. W. Astroski, H. Lin, Y. L. Liu, E. J. Androphy, PloS One 2016, 11.

[22] F. Gros-Louis, R. Lariviere, G. Gowing, S. Laurent, W. Camu, J. P. Bouchard, V. Meininger, G. A. Rouleau, J. P. Julien, J Biol Chem 2004, 279, 45951–45956.

[23] J. M. Beaulieu, M. D. Nguyen, J. P. Julien, J Cell Biol 1999, 147, 531–544.

[24] M. A. Meyer, Neurology International 2014, 6, 5367.

