## Supplemental Information for "Ultrasensitive single-cell proteomics workflow identifies >1000 protein groups per mammalian cell"

b. Thermo Fisher Scientific, San Jose, CA, 95134

c. Biogen Inc, Cambridge, MA, 02142

d. Environmental Molecular Sciences Laboratory, Pacific Northwest National Laboratory, Richland, WA, 99354

### Experiment Section

#### Material and sample preparation

Dithiothreitol (DTT), iodoacetamide (IAA), Pierce Formic Acid (LC-MS grade), and Pierce HeLa Protein Digest Standard were purchased from ThermoFisher Scientific (Waltham, MA). CHROMASOLV™ LC-MS water and acetonitrile were products of Honeywell (Charlotte, NC), MS-grade trypsin and Lys-C were from Promega (Madison, WI). All other chemicals and reagents were purchased from Sigma-Aldrich (St. Louis, MO) unless otherwise noted.

#### Sample collection

HeLa cells (ATCC, Manassas, VA) were cultured and isolated as described previously.[1] Fresh frozen human spinal tissue (ProteoGenex, Los Angeles, CA) was cryosectioned to a thickness of 12 µm and deposited onto PEN-coated microscope slides (Zeiss, Oberkochen, Germany) and fixed with 70% ethanol for 15 min. The tissue sections were stained with hematoxylin and eosin and imaged at 40× resolution using a Zeiss PALM MicroBeam system. Individual motor neurons and interneurons were selected from the ventral horn region and intermediate zone, respectively, of the spinal tissue and excised by laser capture microdissection (LCM) using the PALM MicroBeam system. The excised single neurons were collected into the nanowells of the nanoPOTS chip[2] (Figure S2) and then prepared for analysis as described below.

#### NanoPOTS sample processing

Samples were processed using the nanoPOTS workflow as described previously.[1, 3] Briefly, nanoliter pipetting is accomplished using an in-house-built robotic liquid handling system and a microfabricated glass chip patterned with hydrophilic nanowells arrays. The single cells were collected onto nanowells, and then reagents for cell lysis, reduction, alkylation and digestion were added and incubated sequentially in a one-pot workflow as described previously[3] to generate peptides for analysis.

**LC-MS analysis**

Prior to separation, the sample was transferred from the storage capillary to a home-packed 75-µm-i.d. SPE column for desalting by infusing Mobile Phase A (MP A; 0.1% formic acid in water) at a flow rate of 1 µL/min for 10 min using an UltiMate 3000 RSLCnano pump (Thermo Fisher). The SPE column was then connected to an in-house slurry-packed 20 µm i.d., 50-cm-long LC column[1] with a zero-dead-volume union (Valco, Houston, TX). Chromatographic media for both the SPE and LC columns were 3-µm C18 porous particles having 300 Å pores (Phenomenex, Torrance, CA). The LC separation flow rate was ~20 nL/min, which was split from 250 nL/min programmed flow provided by the same UltiMate 3000 RSLCnano pump using an in-house-prepared 50-cm-long, 75-µm-i.d. column using the same packing material as the analytical column. A linear 100-min gradient of 8-22% mobile phase B (MP B; 0.1% formic acid in acetonitrile) was used for separation. An additional 20-min gradient of 22-45% MP B was used to elute hydrophobic peptides and the gradient was then ramped to 90% MP B over 5 min and held for 5 min to wash the column. Finally, the gradient was ramped to 2% MP B over 5 min and held for 15 min to re-equilibrate the column.

A Thermo Fisher Orbitrap Eclipse Tribrid mass spectrometer with a FAIMS Pro Interface (San Jose, CA) was employed for MS analysis. For data collected without FAIMS, an electrospray potential of 2.0 kV was applied at the source for ionization, while 2200 V was applied when the FAIMS Pro interface was incorporated. The ion transfer tube was set at 150 °C for desolvation and the ion funnel RF level was 30. The Orbitrap mass analyzer was used as the MS1detector with the resolution set at 240,000 (m/z 200).  The resolution was set at 60,000 when the Orbitrap served as the MS2 detector, and the ion trap scan rate was set to Rapid when the ion trap mass analyzer was used for MS2 detection. The MS1 AGC target and maximum injection time were set at 1E6/250 ms, the MS2 AGC target and maximum injection time were set to 1E5/500 ms for the Orbitrap, and 3E4/300 ms when the ion trap was used for detection. Data-dependent acquisition mode was used to trigger precursor isolation and sequencing. Precursor ions with charges of +2 to +7 were isolated for MS2 sequencing. The MS2 isolation window was 1.6 Da, the dynamic exclusion time was set at 120 s, a mass tolerance of ±10 ppm was utilized and the signal intensity threshold was set to 8E3. A normalized collision energy of 30% was used for precursor fragmentation for higher energy collision-induced dissociation (HCD) mode and a normalized collision energy of 35% was used for precursor fragmentation for collision-induced dissociation (CID) mode.

**Data analysis**

The mass spectrometry proteomics data have been deposited to the ProteomeXchange Consortium via the PRIDE partner repository[4] with the dataset identifier PXD019515 (for reviewer access: Username:, Password: a9jWR1UI). Raw files were processed using Proteome Discoverer Software (version 2.4, San Jose, CA) for feature detection, database searching, and protein/peptide quantification. MS/MS spectra were searched against the UniProtKB/Swiss-Prot human database (downloaded on June 6th, 2019, containing 20,353 reviewed sequences). N-terminal protein acetylation and methionine oxidation were selected as variable modifications. Carbamidomethylation of cysteine residues was set as a fixed modification. The mass tolerances of precursors and fragments were <5 and 20 ppm, respectively. The minimum peptide length was six amino acids and the maximum peptide length was 144 amino acids. The allowed missed cleavages for each peptide was 2. A second stage search was activated to identify semi-tryptic peptides. Proteins were filtered with a maximum FDR of 0.01. Both unique and razor peptides were selected for protein quantification. Other unmentioned parameters were the Proteome Discoverer default settings. Potential contaminants from culture media were filtered out using the Bos Taurus Uniprot database. MaxQuant searches performed for comparison used the same search criteria as described previously.[1]

Normalized abundance values for High Confidence (1% protein-level FDR) Master Proteins determined in Proteome Discoverer Software (v2.4) (ThermoFisher Scientific) were loaded into Perseus (v1.6.5.0),[5] log2-transformed and filtered to retain proteins detected in either all three MN samples or all three IN samples. Remaining missing values (~2% of all values) were imputed for the total matrix based on random selection from a normal distribution downshifted by 3 standard deviations (width=0.3 standard deviations). Fold difference in abundance for individual proteins was determined by subtracting the average log2-transformed protein abundance in the IN group (n=3) from the averaged log2-transformed protein abundance in the MN group (n=3). Significance of differential abundance was determined by performing a two-tailed t-test and imposing a significance cutoff threshold of p.adj.<0.05 and ≥2-fold differential abundance on a log2 scale.

Proteins exhibiting significant differences in single MNs vs INs were imported into the web-based STRING (v11)[6] tool for assembly of functional networks allowing a minimum interaction score cutoff of 0.4 and with the text-mining option for active interaction sources disabled. Interaction networks built in STRING were imported into Cytoscape (v3.7.2)[7] to allow mapping of protein abundance data onto individual nodes. P-values for gene ontology (GO) and pathway enrichments were calculated using a Hypergeometric test (statistical background = whole genome) followed by Benjamini-Hochberg correction for multiple hypothesis testing using the STRING enrichment analysis widget.[8]

Supporting Information

Table S1. Protein groups identified from 0.5 ng of HeLa digest

| Method | Protein groups | | | |
|---|---|---|---|---|
| | Replicate 1 | Replicate 2 | Replicate 3 | Average |
| OTOT-HCD 2 FAIMS CVs | 1498 | 1627 | 1633 | 1586 |
| OTIT-CID 2 FAIMS CVs | 1729 | 1844 | 1933 | 1835 |
| OTIT-HCD 2 FAIMS CVs | 2171 | 2009 | 2003 | 2061 |
| OTIT-HCD 3 FAIMS CVs | 1847 | 1937 | 1803 | 1862 |

Table S2. Protein groups identified from single cells and blank sample analyzed by LC/MS

| | Sample | Protein groups | | | | Peptide groups | | | |
|---|---|---|---|---|---|---|---|---|---|
| | | R1 | R2 | R3 | Average | R1 | R2 | R3 | Average |
| FAIMS | SH | 1031 | 1156 | 980 | 1056 | 4064 | 4395 | 3276 | 3912 |
| | SM | 972 | 1192 | 1126 | 1097 | 3063 | 3595 | 3232 | 3297 |
| | SI | 989 | 1265 | 1322 | 1192 | 2978 | 3842 | 4195 | 3672 |
| | Blank | 149 | 200 | 224 | 191 | 320 | 443 | 524 | 429 |
| No FAIMS | SH | 532 | 430 | 418 | 460 | 2460 | 1814 | 1545 | 1940 |

Note: SH: single HeLa cell, SM: single motor neuron, SI: single interneuron; R1,2,3: replicates 1,2,3; Blank: HeLa cell supernatant

Figure S1

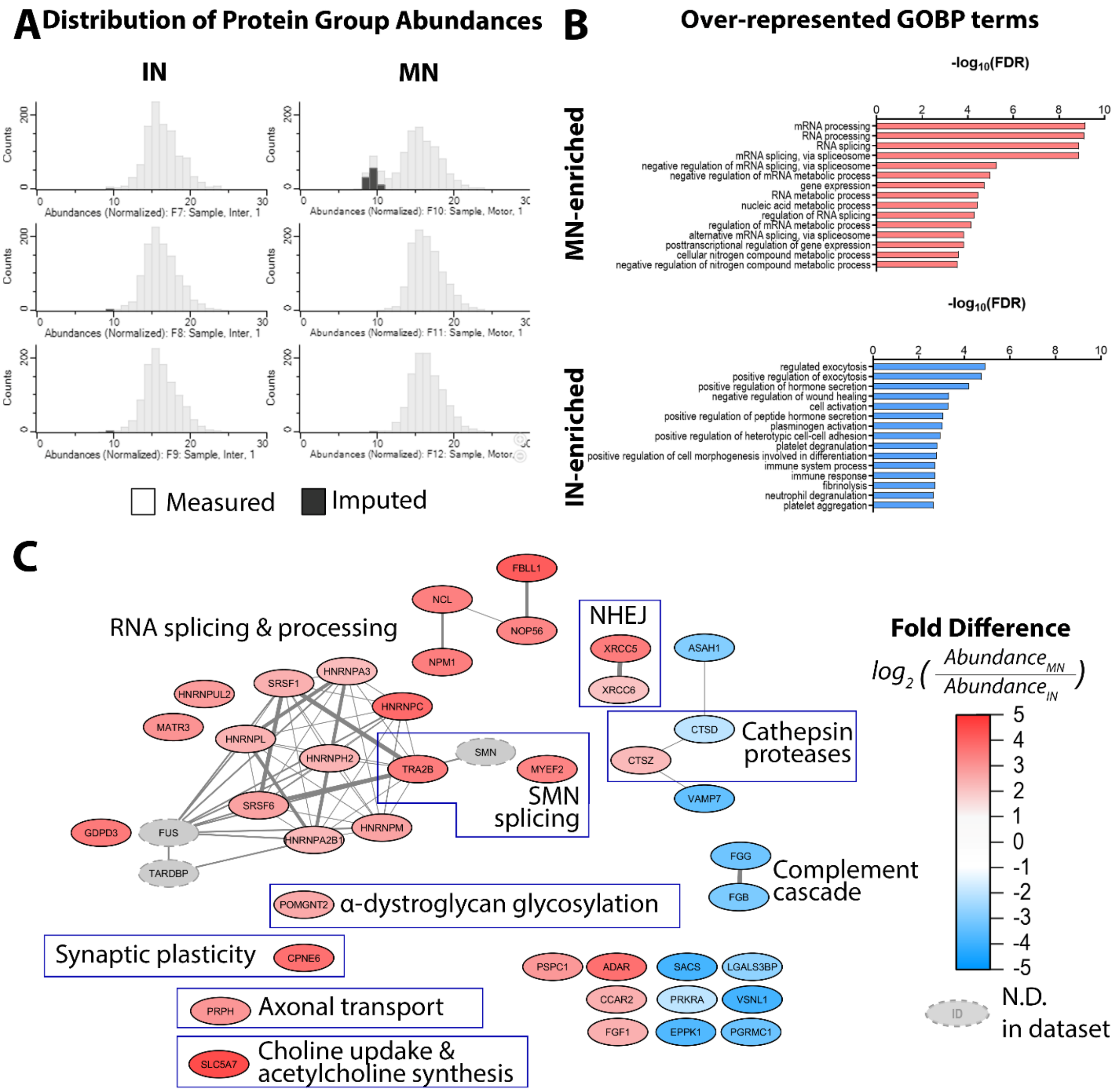


Figure S1. Single-cell proteomic interrogation of human spinal motor neurons (MNs) and interneurons (INs). A) Distribution of log2-transformed values for 1118 quantified protein groups. Imputed values shown in dark gray B) Top 15 significantly (Benjamini-Hochberg-corrected FDR<1%) over-represented Gene Ontology Biological Process (GOBP) categories pathways within the subset of proteins significantly enriched in MNs (28 protein groups, red) or INs (11 protein groups, blue) ($p<0.05$ |Fold Difference|≥2). C) Protein-protein interactions among 39 significantly differentially expressed proteins in MNs and INs. Nodes represent individual protein groups, edges represent protein-protein interactions curated by the STRING database (v11). Node color indicates fold difference in protein group abundance, edge width indicates strength of reported interaction ($0.4<x<1$). Nodes indicated in gray (TARDBP, FUS, SMN) were not detected in single MNs or INs and were manually added to the network as examples of connectivity of enriched-in-MN proteins with protein groups implicated in motor neuron disease.

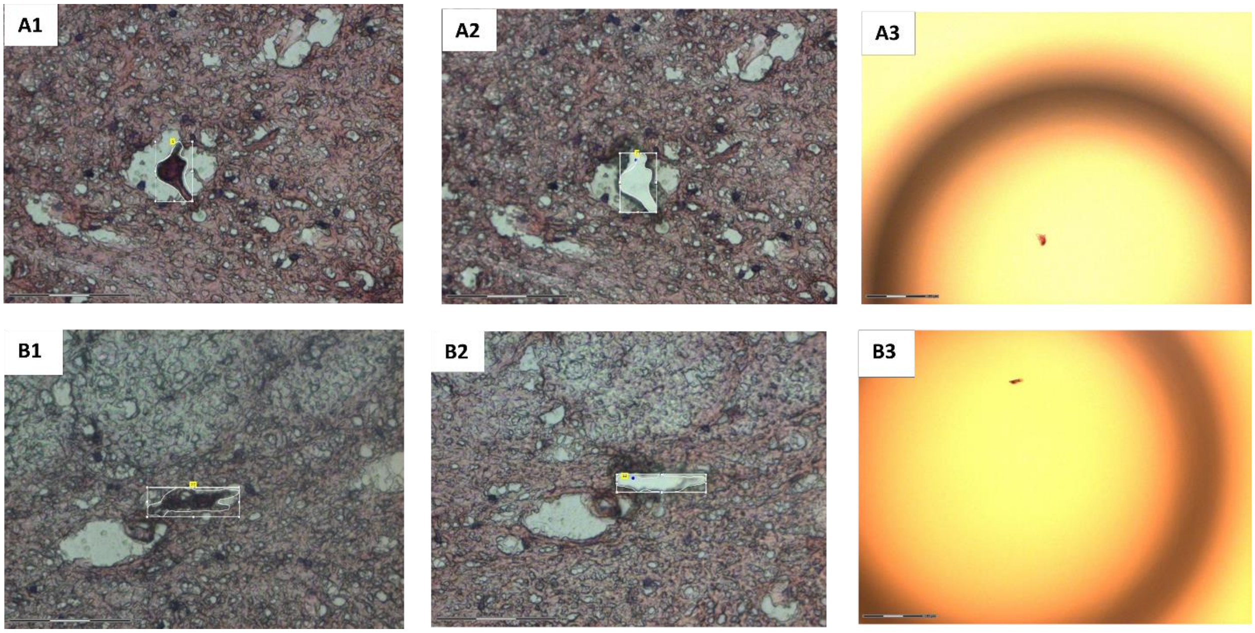


Figure S2. Representative images showing laser capture microdissection of single motor neurons and interneurons. A1 and A2 show H&E stained tissue before and after microdissection of a motor neuron. A3 shows the same motor neuron transferred to a nanowell. B1–B3 show corresponding images for an interneuron.

#### Supplementary References


[1] Y. Cong, Y. Liang, K. Motamedchaboki, R. Huguet, T. Truong, R. Zhao, Y. Shen, D. Lopez-Ferrer, Y. Zhu, R. T. Kelly, *Anal Chem* **2020**, *92*, 2665-2671.

[2] Y. Zhu, M. Dou, P. D. Piehowski, Y. Liang, F. Wang, R. K. Chu, W. B. Chrisler, J. N. Smith, K. C. Schwarz, Y. Shen, A. K. Shukla, R. J. Moore, R. D. Smith, W. J. Qian, R. T. Kelly, *Mol Cell Proteomics* **2018**, *17*, 1864-1874.

[3] aY. Zhu, P. D. Piehowski, R. Zhao, J. Chen, Y. Shen, R. J. Moore, A. K. Shukla, V. A. Petyuk, M. Campbell-Thompson, C. E. Mathews, R. D. Smith, W. J. Qian, R. T. Kelly, *Nat Commun* **2018**, *9*, 882; bY. Zhu, G. Clair, W. B. Chrisler, Y. Shen, R. Zhao, A. K. Shukla, R. J. Moore, R. S. Misra, G. S. Pryhuber, R. D. Smith, C. Ansong, R. T. Kelly, *Angew Chem Int Ed Engl* **2018**, *57*, 12370-12374.

[4] Y. Perez-Riverol, A. Csordas, J. Bai, M. Bernal-Llinares, S. Hewapathirana, D. J. Kundu, A. Inuganti, J. Griss, G. Mayer, M. Eisenacher, E. Perez, J. Uszkoreit, J. Pfeuffer, T. Sachsenberg, S. Yilmaz, S. Tiwary, J. Cox, E. Audain, M. Walzer, A. F. Jarnuczak, T. Ternent, A. Brazma, J. A. Vizcaino, *Nucleic Acids Res* **2019**, *47*, D442-D450.

[5] S. Tyanova, T. Temu, P. Sinitcyn, A. Carlson, M. Y. Hein, T. Geiger, M. Mann, J. Cox, *Nat Methods* **2016**, *13*, 731-740.

[6] D. Szklarczyk, A. L. Gable, D. Lyon, A. Junge, S. Wyder, J. Huerta-Cepas, M. Simonovic, N. T. Doncheva, J. H. Morris, P. Bork, L. J. Jensen, C. V. Mering, *Nucleic Acids Res* **2019**, *47*, D607-D613.

[7] P. Shannon, A. Markiel, O. Ozier, N. S. Baliga, J. T. Wang, D. Ramage, N. Amin, B. Schwikowski, T. Ideker, *Genome Res* **2003**, *13*, 2498-2504.

[8] A. Franceschini, D. Szklarczyk, S. Frankild, M. Kuhn, M. Simonovic, A. Roth, J. Lin, P. Minguez, P. Bork, C. von Mering, L. J. Jensen, *Nucleic Acids Res* **2013**, *41*, D808-815.
